# Protein trafficking and synaptic demand configure complex and dynamic synaptome architectures of individual neurons

**DOI:** 10.1101/2025.07.31.667853

**Authors:** Oksana Sorokina, Edita Bulovaite, Anatoly Sorokin, Seth G.N. Grant, J. Douglas Armstrong

## Abstract

Excitatory synapses are the most abundant synapse type in the brain. Being essential for behaviour and are implicated in hundreds of brain disorders, these synapses exhibit striking structural and functional diversity. Synaptome mapping at single-synapse resolution reveals that synaptic protein diversity is spatially organised along the dendritic tree of individual neurons and varies with age and cell-type. However, the cell biological mechanisms underlying the generation of these complex spatial synaptic patterns remain poorly understood. Potential mechanisms include somatic and dendritic protein synthesis, protein trafficking, and local regulatory mechanisms such as activity-dependent degradation. Here we developed computational models to test how combinations of these processes account for empirical synaptome data. We found that a combination of molecular transport mechanisms and local synaptic demand for proteins was sufficient to explain very complex profiles of synaptic protein distributions observed in young, mature and old mice and in different cell types. Our findings suggest the highly complex and dynamic synaptome architecture of the brain is an emergent property of a minimal set of cell biological processes. Our model sets the stage for simulations of brain tissue incorporating molecularly diverse neuronal and synaptic types in a synaptome and connectome architecture.

## Introduction

The hallmark anatomical feature of neurons is their highly elaborate tree-like branches that maximise the number of synaptic contacts^1^. A single neuron may receive direct input on its dendritic tree from thousands of other neurons. Every synaptic contact contains up to hundreds of thousands of individual proteins encoded by several thousand genes^2^. These proteins assemble into multiprotein complexes^3^ that are molecular machines mediating release of neurotransmitters from the presynaptic terminal and signal integration in the postsynaptic terminal. The crucial role of these proteins is exemplified by the hundreds of gene mutations that result in brain and behavioural disorders^4,5^.

The majority of synapses in the mammalian brain employ glutamate as a neurotransmitter and are excitatory in nature. However, underneath the common transmitter phenotype there is a hidden degree of complexity at the individual synapse level with vastly diverse protein compositions. The proteomes of individual synapses differ depending on their location on the dendritic tree, neuron type, age, neuronal activity, genetics and other factors ^6–8^. Additionally, synapse proteins have to be replaced continuously and the time course of their turnover differs between synapses, further contributing to the molecular diversity of synapses ^9,10^.

The spatial distribution of molecularly diverse synapses on the dendritic tree is non-random. For example, synaptome mapping reveals the synaptic scaffolding proteins PSD95 and SAP102^6^ are differentially distributed on the dendrites of CA1 pyramidal neurons (PyNs) and dentate gyrus granule neurons (DGs) in the intact mouse brain. Single-synapse resolution mapping of PSD95 turnover also indicates the presence of complex spatial gradients across the dendritic tree ^9^. However, these gradients are not simple continuums from soma to distal dendrite but appear to be a combination of such continuums together with local discontinuities that correlate with the presence of pathway-specific axonal inputs ^9^. This suggests the interplay of two mechanisms: a cell-autonomous mechanism that establishes a distance-dependent gradient and a non-cell autonomous mechanism that operates in local regions of the dendrite ^6,7,9^.

To explore the potential mechanisms underlying complex gradients of diverse synapses in neurons, we adapted existing computational models of dendritic molecule distributions to the synaptome mapping data that reveals these gradients in mice of different ages. An extensive literature^11^ invokes many mechanisms including somatic and dendritic protein synthesis, active and passive trafficking of proteins to their destination and local protein degradation. While simulations incorporating diffusion, active transport and local translation have been explored, none have been applied to the complex molecular and synaptic gradients observed in the intact brain^11^.

Here we used a model that incorporates Doyle and Kiebler’s ‘sushi-belt’ theory of molecular transport^12,13^, which assumes that molecules synthesised in the soma are continuously trafficked throughout the entire dendritic tree, imitating a conveyor belt. Local demand for proteins, which mimics the activity/plasticity state of the local synapse population, generates hot spots of “hungry” synapses that sequester and anchor these molecules. This model can be conveniently simulated with the neuron geometry defined in a format compatible with NEURON, a biophysical multi-compartment model^13,14^, accommodating the intricate branching of the neuron type in question.

We showed that the complex gradients of synapse composition and dynamics can be simulated in two major classes of excitatory neurons – pyramidal and granule cell neurons – using a model assuming only somatic protein synthesis and a sushi belt that distributes proteins to synapses with different demand and degradation rates. Our simulation recapitulates the synaptome architecture observed in mice of different ages across the lifespan, implicating the regulation of specific cellular processes that change with age. The arising picture is that the spatial organisation of molecular diverse synapses could be an emergent property of a few basic cell biological processes.

## Results

### Mapping the spatiotemporal distribution of PSD95 in hippocampal pyramidal neurons onto the computational model

Bulovaite et al, 2022^9^ measured the quantity of fluorescently pulse-labelled synaptic PSD95 puncta (density; puncta number per unit area) in 20 individual subregions of CA1 pyramidal neurons in the mouse brain and tracked the labelled protein over seven days. These data revealed complex distributions of synapse densities in neurons. Further, they showed a complex dynamic process where the amount of labelled PSD95 (intensity of the fluorescent signal for each puncta) at different regions of the neuron changed over several days. As a result of the progressive loss of PSD95, the synaptic puncta intensity diminishes as does the number of visible synapses (density). Figure 1A shows the distribution of synapses with different densities at day 0. In apical dendrites, the lifetime of PSD95, hereafter referred to as synapse protein lifetime, appeared to increase with distance from the soma. However, the curve was not linear and included a dip at the intersection between stratum radiatum and stratum lacunosum of the CA1. A gradient of increasing synapse protein lifetime with distance from soma was also observed in the more proximal parts of stratum oriens before levelling and reversing in the most distal regions.

**Figure 1.**
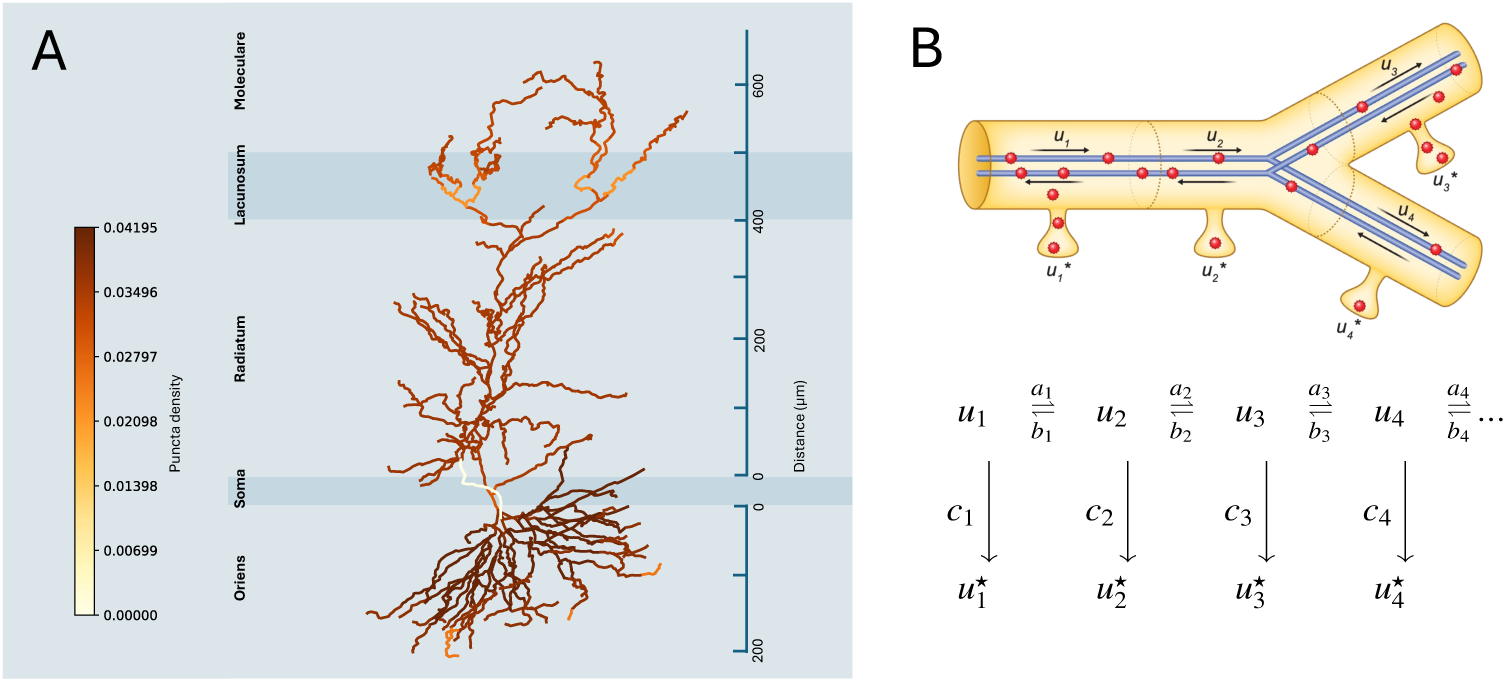
A. Model of a pyramidal CA1 neuron colour coded by values for normalised PSD95 puncta density taken on day 0 of the Halo-ligand injection (from ^15^). The apical dendrite traverses three subregions of the hippocampus (stratum radiatum, lacunosum and moleculare) and the basal dendrites occupy the stratum oriens. Each subregion of the CA1 receives distinct afferent inputs: CA1 stratum radiatum receives projections from CA3, while stratum lacunosum is targeted by entorhinal stellate cells in layer 2 and stratum moleculare by entorhinal cortex layer 3. Stratum oriens receives inputs from CA2 and CA3 pyramidal neurons. B. Upper: schematic representation of four neighbouring sushi-belt model compartments, with cargo (u1, u2, u3, u4) and synaptic (u1*, u2*, u3*, u4*) PSD95 in each compartment. Compartments are enumerated from soma to most distant part to dendritic tree. Lower; mass-action model with respective kinetic constants taken from ^13^.

The synapse protein lifetime was measured in a total of 20 compartments assigned to regions of the dendritic tree of PyNs^15^. PSD95 puncta density was quantified across a series of windows starting from the soma of the pyramidal neuron in CA1 through to the distal basal dendrites in stratum oriens (so) and to the distal apical dendrites in stratum radiatum (sr), stratum lacunosum (sl), and stratum moleculare (sm) (Figure 1, A). Each of these dendritic regions were subdivided into several compartments: 5 in so, 10 in sr and 5 in sl and sm together (slm in Figure 1, B) giving a total of 20 subregions. A schematic representation of the CA1 neuron structure is presented in Figure 1 (A, upper panel and B, left panel).

In common with previously described models for molecular trafficking in CA1 neurons ^16,13^ and which are discussed further in the next section, we considered the CA1 neuron to consist of 742 compartments spanning five types. Given that we do not have PSD95 values for the soma or axon, we only considered the “dend”, “dend5” and “apic” compartment types. Those three types were mapped onto the experimental data using their relative distance from the soma (Table 1). Given the highly branched structure of the neuron (Figure 1, B, right panel), compartments the same distance from the soma all inherit the same value for puncta density. Code for mapping is available as a part of Jupyter notebook on GitHub.

**Table 1.** Mapping of CA1 neuron subregions to the model of reconstructed pyramidal neuron. Abbreviations for CA1 neuron subregions mapped on the model compartments from ^16^. Each of 20 subregions measured in experiment correspond to 6-134 model compartments located at the equal distance from the soma measured along the vertical/Z-axes).

| Neuron subregion number | Neuron subregion name | Subregion abbreviation | Distance from the soma (along Z-axes) | Model compartment name | Number of model compartments |
| --- | --- | --- | --- | --- | --- |
| 1 | CA1so | CA1so_1 | -165 | dend | 9 |
| 2 | CA1so | CA1so_2 | -127.5 | dend | 85 |
| 3 | CA1so | CA1so_3 | -90 | dend | 134 |
| 4 | CA1so | CA1so_4 | -52.5 | dend | 100 |
| 5 | CA1so | CA1so_5 | 0 | dend | 16 |
| 6 | CA1sr | CA1sr_1 | 10 | apic | 3 |
| 7 | CA1sr | CA1sr_2 | 52.5 | apic | 6 |
| 8 | CA1sr | CA1sr_3 | 94.286 | apic | 29 |
| 9 | CA1sr | CA1sr_4 | 136.43 | apic | 37 |
| 10 | CA1sr | CA1sr_5 | 178.57 | apic | 48 |
| 11 | CA1sr | CA1sr_6 | 220.71 | apic | 44 |
| 12 | CA1sr | CA1sr_7 | 262.86 | apic | 45 |
| 13 | CA1sr | CA1sr_8 | 305 | apic | 38 |
| 14 | CA1sr | CA1sr_9 | 347.14 | apic | 21 |
| 15 | CA1sr | CA1sr_9 | 389.29 | apic | 9 |
| 16 | CA1slm | CA1slm_1 | 431.43 | dend5 | 13 |
| 17 | CA1slm | CA1slm_2 | 473.57 | dend5 | 24 |
| 18 | CA1slm | CA1slm_3 | 515.71 | dend5 | 35 |
| 19 | CA1slm | CA1slm_4 | 557.86 | dend5 | 29 |
| 20 | CA1slm | CA1slm_5 | 600 | dend5 | 9 |

Assuming the total concentration of (labelled and unlabelled) PSD95 to be more or less constant in any given puncta, the synapse protein lifetime of labelled synaptic PSD95 across the subcellular regions may be a compound of at least two processes: First, there is the actual protein turn-over, or molecular decay rate, where labelled protein is degraded and subsequently replaced by new unlabelled protein. Second, there will be a rate of replenishment of degraded protein with pre-existing labelled PSD95 from the finite pool existing elsewhere, including from molecular transport mechanisms such as a sushi-belt conveyor. We asked if variations in the rate of these processes within dendritic compartments could account for the observed spatial differences in synapse protein lifetime.

### Extending the sushi-belt model for protein distribution simulations

Williams et al’s (2016)^13^ model (Figure 1, B) was designed to predict molecular distribution given a set of parameters, based on an assumption that the molecule pool moves from soma to the points of high local demand. Here, we face the reverse challenge; can we calculate the model parameters that best fit the observed molecular distributions? Therefore, we implemented a new model, using the same underlying mathematical principles, but enabling simulations that can be used to explore the parameter space and allow us to compare predicted profiles against observed data (further details in Methods).

In common with Williams et al (2016), our PSD95 cargo is bound to motor proteins, which can move the cargo in both directions along the dendritic branches. The model splits these branches into virtual compartments and considers cargo movement between adjoining compartments. The reconstructed 3D pyramidal neuron model ^16^ is divided into 742 compartments of varying length from 5 to 34 microns. Therefore, as a first step, we needed to align them to the 20 subregions from the original experimental data, where each compartment in the model inherits a corresponding value for the PSD95 puncta density (Table 1) based on their distance from the soma (see also Figure 1 and Methods).

The parameter *u* represents the concentration of PSD95 ‘cargo’ attached to the microtubules and is indexed by compartment. PSD95 can move to neighbouring compartments with rate constants *a* or *b,* for forwards or backwards movement, respectively. Cargo can also irreversibly detach from the microtubule with detachment constant *c,* thus providing a concentration of detached cargo *u*,* which – given the nature of the data – we assume to be mostly ‘synaptic’ (Figure 1, B).

As was suggested in ^13^, the interplay between the different transport processes available in the model result in behaviours that lie on a spectrum between two extreme modes: One end is dominated by demand-dependent traffic (DDT), where the rate of movement depends on the local demand and the rate of detachment of cargo from the transport system is equal across all compartments. At the other end, demand-dependent detachment (DDD), where the rate of movement is uniform across the cell, while the rate of detachment varies with local demand. The ratio of these modes is represented by the parameter *F* (see Appendix).

In contrast to the Williams model, where all cargo starts at the soma, we already have fully distributed cargo at the outset. Therefore, we extended the model by adding an additional parameter *mProp*, which defines the ratio of cargo:synaptic PSD95 at simulation time 0. Moreover, the original model does not consider degradation rates at synapses, which is clearly a fundamental process affecting the time course of the experimental dataset. Therefore, we further extended the model to add another parameter *d* describing the molecular degradation rate for synaptic PSD95 protein. This extended model can simulate the impact of varying rates of trafficking, detachment (*a*,*b* and *c*) and degradation *d,* to assess how well these can account for the distribution of protein observed in the biologically obtained datasets.

### Inferring the local demand profile from protein distribution data

Our simulations are based on a series of assumptions as follows: First, we assume that puncta density, as measured by ^9^, correlates with the underlying local demand for PSD95 at the location of the signal. Thus, we can estimate local demand values from fitting the sushi-belt model to the PSD95 puncta density profile. Second, we assume that protein bound to the polymerized microtubule bundles (*i.e*. cargo) is not measured as it forms the background of the original imaging data. Therefore the measured puncta represent visible pools of PSD95 which we assume to be mostly synaptic. Moreover, in the experimental data, only proteins expressed during day 0 are available to the HaloTag ligand and can subsequently be measured by imaging. Therefore, we can safely ignore the influence of local and somatic translation mechanisms in explaining observed changes in distribution of labelled PSD95 protein between day 0 and day 7. Next, we assume that the neuron is, on average, in a steady state across the 7 days; the total PSD95 at each synapse remains relatively constant but the ratios of labelled vs unlabelled protein change over the time course resulting in the values for synapse protein lifetime. For simplicity, we assume that molecular degradation occurs only at the detached protein (*e.g*. synaptic) and does not affect cargo protein in the active transport system. While our model can support reattachment of synaptic protein back onto the cargo pool, again for simplicity we have excluded this from simulations.

### Spatiotemporal distribution of PSD95 is tuned by local transport

We tested how well the model captures the observed changes in synapse protein lifetime of PSD95 protein across the 20 dendritic regions. We used the Day 0 data to indicate the initial abundance of synaptic PSD95. We then simulated the distribution of synaptically localised protein at Day 7 by adjusting the parameters for traffic, detachment and degradation. Finally, we compared the obtained protein distribution with experimental values (after normalisation for non-uniform number of compartments and Day 0 distribution, for details of the optimisation see Methods).

Given the original complexity of the ^16^ model (742 compartments), we tried to minimise model overfitting by keeping the parameter number as low as possible. We started from a model with just three parameters for demand, assuming that demand values might be uniform within each of the three major CA1 regions: CA1so, CA1sr and CA1slm (Figure 1, B). For this simulation, the degradation rate was set to a constant for the whole cell. In addition to the three local demand values and a global degradation rate *d,* we also fit detachment rate *Ctau*, *F* (ratio between DDD and DDT modes) and *mProp* (ratio between bound and unbound protein at time 0), resulting in seven parameters in total.

We assess each iteration of model for how well the best fit configuration matches the relevant biological dataset (effectively mean squared difference, see Methods). The best fit using this first configuration (Figure 2, A; cost = 0.00024 orange simulation line) does not precisely follow the experimental day 7 data (blue): Deviations are more obvious in the more distant subregions of apical dendrites, where the trends of experimental and simulation curves critically diverge. Therefore, we concluded that the three regions demand model with uniform degradation is not fully sufficient to describe the experimental data. The optimal value for *F* obtained in this simulation is 0, corresponding to a DDD mode, which means that the model achieves the best fit to observed protein distribution by adjusting the detachment rate while keeping traffic rates uniform.

**Figure 2.**
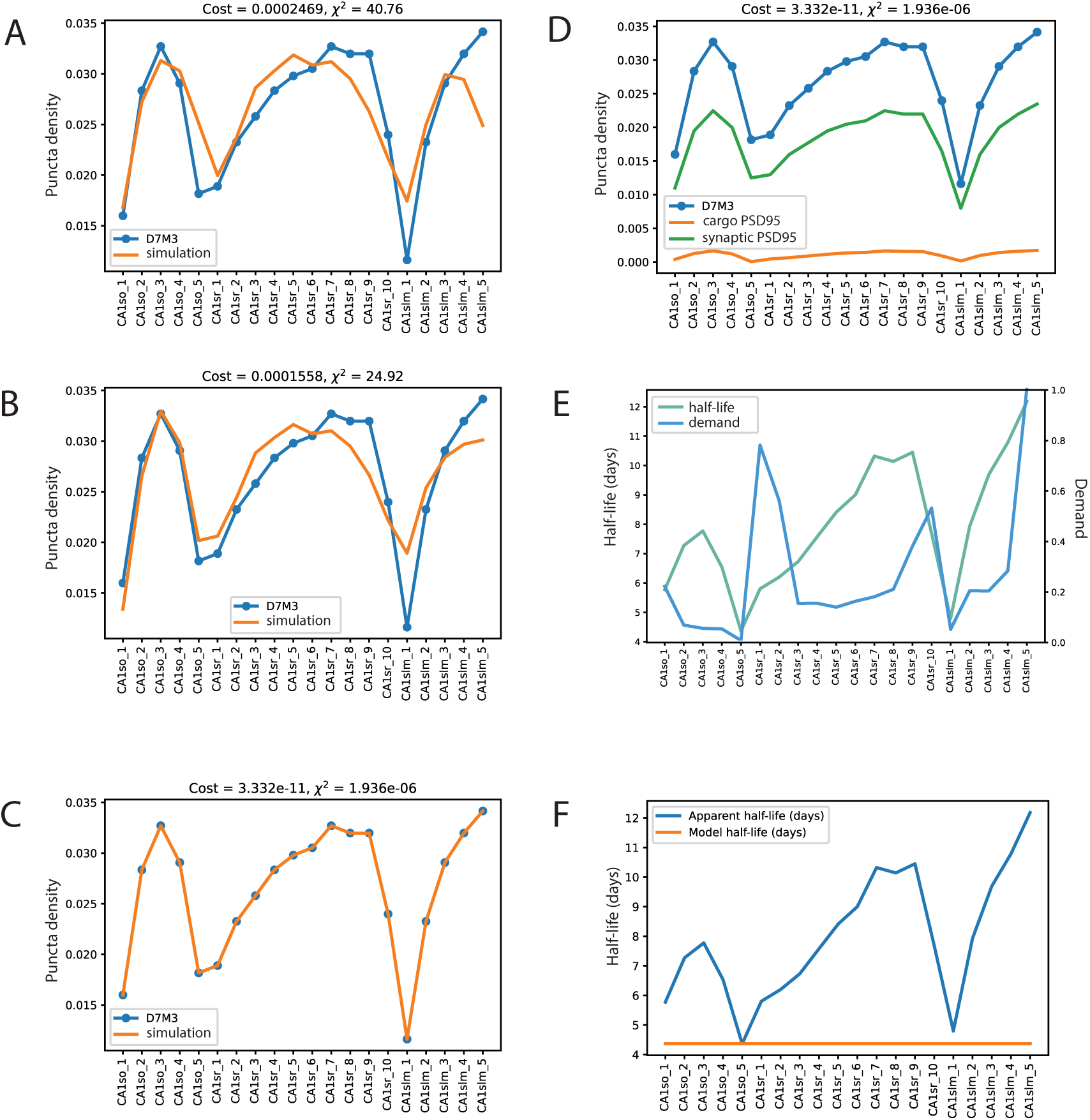
Fitting of the sushi-belt model to the experimental data from three month old mice (error bars removed for clarity, cf Fig 4); D7M3 is a normalized protein distribution obtained from Day 7 puncta density data. A. Best fit for a simple model with three compartments and one degradation rate (blue line with dots – experimental data, orange line – simulation) B. Fitting of model with three compartments and three degradation rates. C Fitting results of final model of 20 compartments and two parameters for degradation rates (blue and orange lines are congruent). D. for the best fit as presented in C shown the both the synaptic protein pool obtained from model (green, not normalised, see Methods for details) and cargo / belt-bound protein pool (orange, 30 % of total as corresponds to mProp = 0.3). E. Apparent half-life estimated from experimental data based on an assumption of exponential decay (green line) is plotted against estimated local demand for each compartment (blue line). F. Apparent half-life estimated from experimental data (blue line) plotted against model half-life (orange line) estimated from simulations.

Next, in the second configuration, we added three distinct degradation rates, one for each of the regions, with its own demand value thus increasing the number of parameters to nine. As shown in Figure 2, B the fit of this model is more accurate (*cost* = 0.00015). The trends for measured and modelled protein distributions at day 7 are now more similar across the proximal regions of the neuron. However, the model still fails to describe the most distant apical dendrite subregions accurately. The obtained value for *F* in this second set is close to 1 (0.99), which means that DDT (demand-dependent traffic) becomes now the dominant process. We noted that in this set of simulations we found that degradation rates correlate positively with demand.

In the final (third) set of simulations, we increased the number of parameters for demand, making it equal to the number of subregions (20) in the experimental data where each of the subregions has its own characteristic value for PSD95 puncta density and synapse protein lifetime. To limit the number of parameters needed we did not implement 20 individual degradation rates, rather we used a linear model with two parameters based on our second simulation results, that degradation is positively correlated with demand:

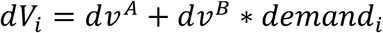

Here, *demand_i_* is the demand value for a specific compartment *i*. *dv^A^* and *dv^B^* are the rates for degradation with *dv^A^* a constant degradation component across the whole neuron and *dv^B^_i_* represents a subregion specific degradation component that correlates with local demand. Note that using a single dV value did not perform well (data not shown).

With this increase in complexity (Figure 2, C), the optimised fit perfectly matches the experimental curve (the orange and blue curves are effectively overlaid). The resulting parameter values are presented in Table 2, with *F* now taking a value of 0.8, which indicates that the system operates primarily in DDT mode, so that redistribution of proteins is achieved mainly by tuning local transportation rate. Meanwhile, the value for *mProp* is 0.3, corresponding to a distribution of 30% of cargo PSD95 and 70% synaptic (Figure 2, D).

**Table 2.** List of parameters obtained in the optimisation for 3-month, 3-week and 18-month old mice in CA1 neurons. Bold indicates most striking changes between models.

| Parameter name | Definition | Fitted value<br>3m | Fitted<br>value 3w | Fitted<br>value<br>18m |
| --- | --- | --- | --- | --- |
| F | Ratio between DDT (1) and DDD (0) modes | 0.8861 | 0.8861 | 0.8861 |
| Ctau | Coefficient for demand-dependent detachment | 4.38e-06 | 4.38e-06 | 4.38e-06 |
| mProp | Ratio between mobile and detached polls protein pools | 0.312 | 0.312 | 0.312 |
| dvA | Intercept for degradation linear approximation (1/s) | 1.837e-06 | **5.21e-06** | 1.63e-06 |
| dvB | Slope for degradation linear approximation (1/s) | 1.002e-18 | 1e-18 | **9.57e-08** |
| Demand_CA1so_1 | Local demand value for st.oriens subregion 1 | 0.214 | 0.214 | 0.214 |
| Demand_CA1so_2 | Local demand value for st.oriens subregion 2 | 0.061 | 0.061 | 0.061 |
| Demand_CA1so_3 | Local demand value for st.oriens subregion 3 | 0.047 | 0.047 | 0.047 |
| Demand_CA1so_4 | Local demand value for st.oriens subregion 4 | 0.045 | 0.045 | 0.045 |
| Demand_CA1so_5 | Local demand value st.oriens subregion 5 | 1.0e-07 | 1.0e-07 | 1.0e-07 |
| Demand_CA1sr_1 | Local demand value for st.radiatum subregion 1 | 0.773 | 0.773 | 0.773 |
| Demand_CA1sr_2 | Local demand value for st.radiatum subregion 2 | 0.554 | 0.554 | 0.554 |
| Demand_CA1sr_3 | Local demand value for st.radiatum subregion 3 | 0.147 | 0.147 | 0.147 |
| Demand_CA1sr_4 | Local demand value for st.radiatum subregion 4 | 0.147 | 0.147 | 0.147 |
| Demand_CA1sr_5 | Local demand value for st.radiatum subregion 5 | 0.131 | 0.131 | 0.131 |
| Demand_CA1sr_6 | Local demand value for st.radiatum subregion 6 | 0.154 | 0.154 | 0.154 |
| Demand_CA1sr_7 | Local demand value for st.radiatum subregion 7 | 0.172 | 0.172 | 0.172 |
| Demand_CA1sr_8 | Local demand value for st.radiatum subregion 8 | 0.202 | 0.202 | 0.202 |
| Demand_CA1sr_9 | Local demand value for st.radiatum subregion 9 | 0.373 | 0.373 | 0.373 |
| Demand_CA1sr_10 | Local demand value for st.radiatum subregion 10 | 0.524 | 0.524 | 0.524 |
| Demand_CA1slm_1 | Local demand value for st.lacunose and molecular subregion 1 | 0.043 | 0.043 | 0.043 |
| Demand_CA1slm_2 | Local demand value for st.lacunose and molecular subregion 2 | 0.196 | 0.196 | 0.196 |
| Demand_CA1slm_3 | Local demand value for st.lacunose and molecular subregion 3 | 0.196 | 0.196 | 0.196 |
| Demand_CA1slm_4 | Local demand value for st.lacunose and molecular subregion 4 | 0.275 | 0.275 | 0.275 |
| Demand_CA1slm_5 | Local demand value for st.lacunose and molecular subregion 5 | 0.999 | 0.999 | 0.999 |

### Preferential transport of PSD95 to the distal ends of the dendritic tree

When we estimate the global degradation rate (*dv^A^*) from the optimised model, we get a value corresponding to a half-life of 4.365 days; which aligns well with other experimentally measured values for PSD95 protein (*e.g*. Cohen et al., 2013, etc)^17^. The value for the distance/subregion dependant component *dv^B^* (1.0e-18) is very small, which suggests that the differences in molecular degradation rate across the compartments are likely to be minimal.

Bulovaite et al., (2022)^9^ reported an “apparent” protein half-life directly from the experimental imaging data (assuming that the change in concentrations between Day7 and Day 0 fits an exponential decay function within each subregion and that there is no influx of labelled protein). The apparent half-life follows a gradient of increasing PSD95 protein half-life observed in stratum radiatum, followed by significant reduction in stratum lacunosum and then, again an increase towards the most distant subregions of stratum moleculare, (see Figures 2E green line & 2F blue line, Supplementary figure 1).

Given that our model indicates a flat distribution for protein half-life (Figure 2, F, orange line) and that the values for local demand do not closely follow the apparent protein half-life reported, we asked what else could account for the redistribution of dendritically localised fluorescently labelled PSD95 protein between Day 0 and Day 7. From a correlation dendrogram of parameter values (Figure 3, A), we can see that the profile of apparent half-life (t_1/2_) correlates with the parameters *a* (forward traffic rate constant) and *c* (detachment rate), which suggests that the labelled PSD95 distribution can be explained by tuning the movement of bound protein towards the distal dendritic compartment, its sequestration by synapses, followed by degradation. See also Supplementary figures 2 and 3.

**Figure 3.**
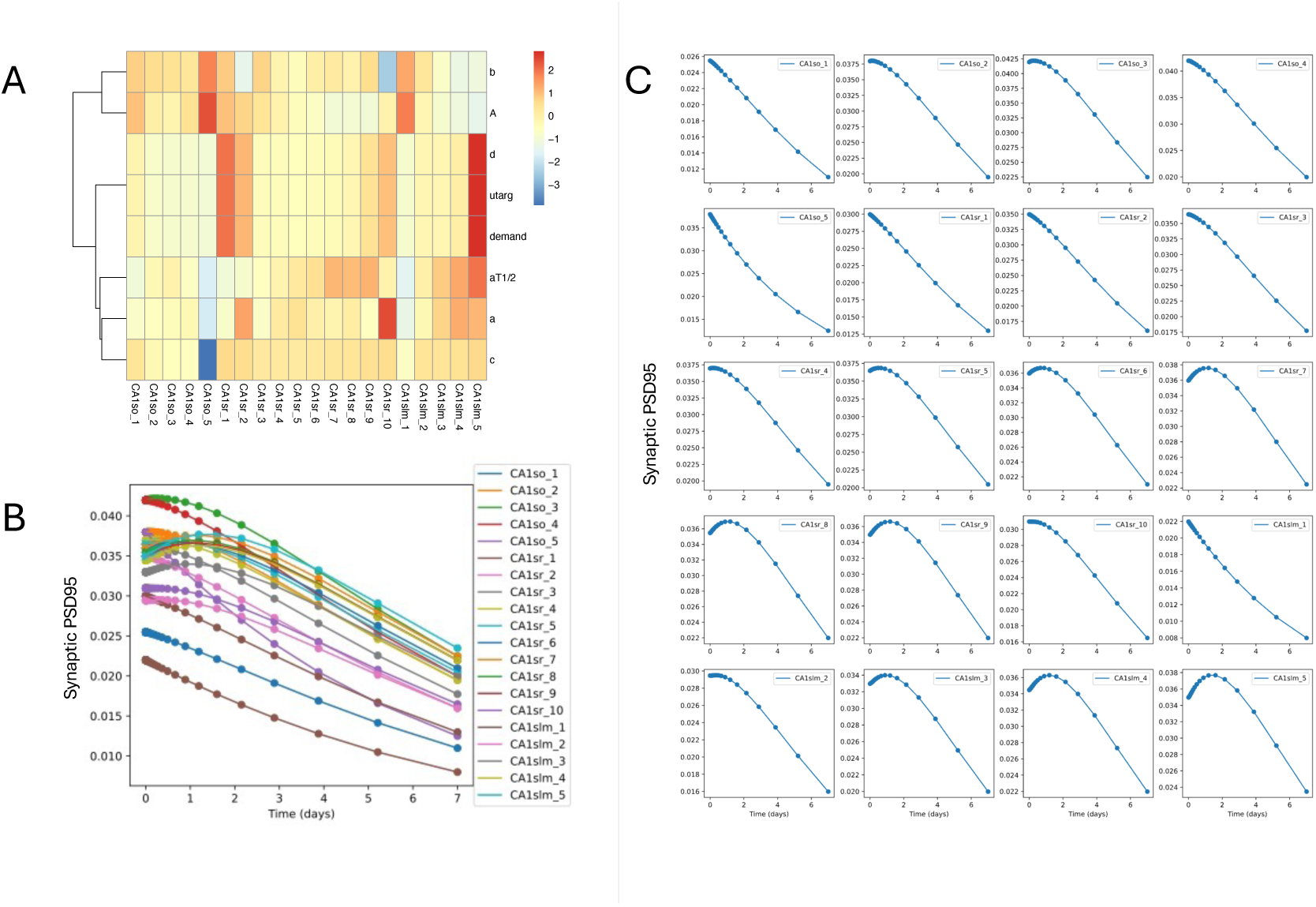
Results of model optimisation. A. Heat map of z-normalized parameters (shown as row names), which are dependent on local demand, including the apparent half-life and the local demand itself. B. Time series of changes in synaptic PSD95 abundance estimated for each compartment based on model prediction C. Similarly to B, but changes in synaptic PSD95 are plotted individually. For CA1sr1 through CA1sr9, peak PSD95 concentration occurs later in more distal compartments which suggests that the protein is moving towards the distal compartments.

Figure 3, B and C show how the predicted concentration of labelled PSD95 protein changes in each of the considered neuron subregions from Day 0 to Day 7. The only two compartments where the amount of PSD95 decays exponentially is CA1so_5 and CA1slm_1, and there, the apparent and model half-lives are similar to each other (Figure 2, F). In other compartments, the amount of protein increases from Day 0 to Day 2 due to transport and then decreases by Day 7. The highest increase at Day 2 can be seen in subregions with the highest apparent half-life, such as CA1sr_7-9, and CA1slm_3-5.

### PSD95 protein half-life varies with age

We tested if the same model parameters could also predict the distribution of PSD95 protein in mice of different ages. According to the data from ^9^, the apparent half-life of PSD95 increases from that seen in juveniles (3-week old mice) to older adults (18-month old mice). We took the best fit model (for 3-month data) and its associated parameters and applied it directly to the data from 3-week and 18-month old animals (Figure 4, A and B), again using their respective puncta density values as initial conditions.

**Figure 4.**
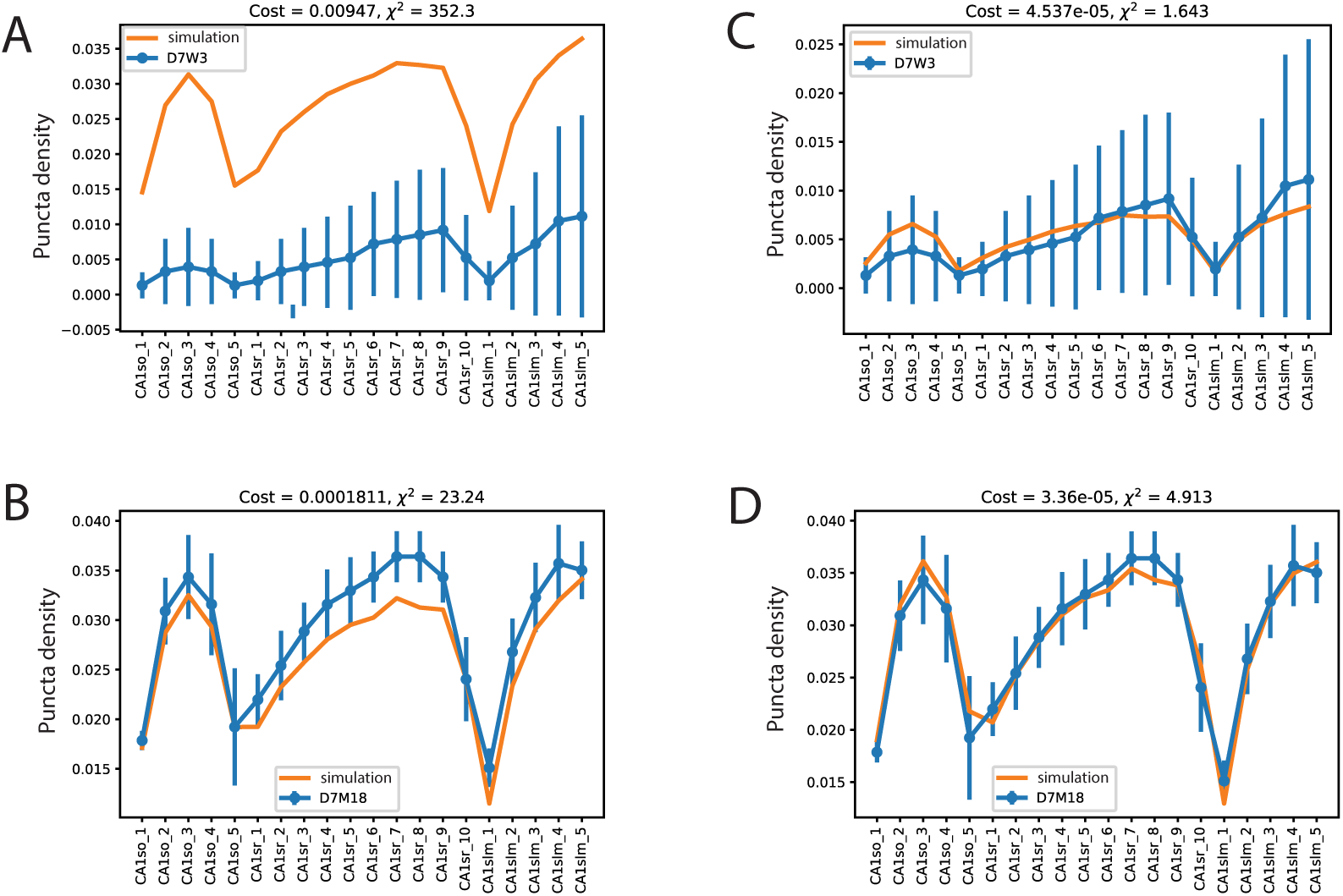
Modelling PSD95 in mice younger (3 week) and older (18 months). Error bars represent experimental SD from ^15^. A. D7W3 is the normalized protein distribution obtained from 3-week old mice 7 days post labelling (blue) against the simulation (orange) using the best parameters from the 3-month old simulations. B. As Panel A but using data from 18-month old mice. C. As panel A (3-week old) except with a re-optimised degradation rate. D. As B (18-month old) except with a re-optimised degradation rate.

The model predicts a distribution for 3-week old animals that has a similar shape/profile. However, it overestimates the abundance of PSD95 by a considerable margin (Figure 4, A orange model vs blue data). At 18-months, the model prediction is actually very close to the observed data (Figure 4, B). The similar profiles indicate that the core processes are relatively stable with age. The most likely explanation for the difference at 3 weeks would be a faster molecular degradation rate. Therefore, we adjusted the model to see if allowing degradation rates *dv^A^* and *dv^B^* to vary would allow a better fit. In the 3-week data, a change in *dv^A^* from 1.837e-06 to 5.41e-06 allows an optimal fit and suggests a change in protein half-life between 1.54 days at 3 weeks old compared to 4.3 days observed in mature animals. For a best fit at 18 months, *dv^A^* remains largely unchanged while *dv^B^* has a modest increase from 1.002e-18 to 1.0e-08, indicating that that while global cell rate of degradation rate is constant, there is a trend tends towards degradation being slightly more sensitive to local demand.

### The sushi-belt model can explain PSD95 distribution in other types of neurons

We tested if the model was sufficiently generic to be relevant to other neuron types/architectures. Specifically, we focused on the distribution of PSD95 protein in dentate gyrus (DG) granule cells. In contrast to CA1 PyNs, there are few obvious boundaries in DG cells. Here, the PSD95 imaging data suggests a small linear increase in synapse protein lifetime scaling with distance from soma (with the exception of the most distal regions). The linear increase was barely visible in 3-week old animals.

The experimental data was divided into 10 sections of even width from the soma (Figure 5, A). The model we selected (Neuromorphe ID: NMO_80550), reconstructed from the DG of 2 month-old mice^18^ was divided into 32 compartments (which equalled the number of branches). To make the DG and CA1 PyN models more comparable, we increased the number of compartments, in proportion to the branch length, to 287. As with the CA1 PyN model, these compartments inherited the experimental values based on the data from equivalent distances from the soma.

**Figure 5.**
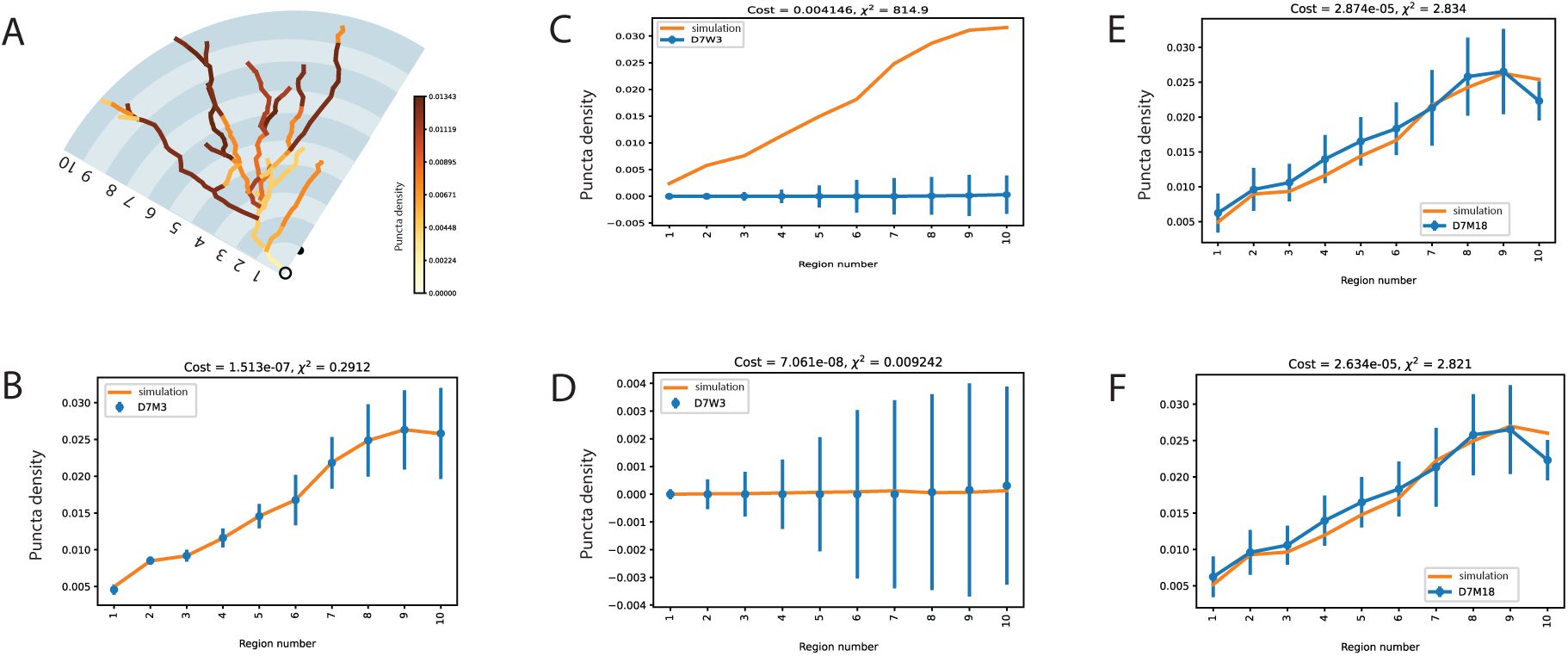
Sushi-belt model application to the PSD95 distribution in Dentate Gyrus cell. A. Schematic image of granule cell obtained from Neuromorpho model (ID: NMO_80550). Panel A shows the 10 subregions considered in the cell. Colour coding indicates normalised puncta density measured in 3-month old mice seven days after labelling. B. Result of fitting the sushi-belt model (orange line) to the experimental data (blue line) of 3-month old mice. C. As B, except comparing a 3-month simulation to data from 3-week old animals. D. As C, except allowing optimisation of the degradation rate. E. As B, except comparing 3-month simulation to data from 18-month old mice. F. As E, except allowing optimisation of the degradation rate.

Given the relatively simpler cell morphology compared to CA1 PyNs together with a basic gradient of synaptic puncta density distribution, we decided to start from a simpler model of 10 subregions with 10 local demand values and one degradation rate. Figure 5, B shows this model already fits the experimental data (*cost* = 1.51e-07) and did not require any increase in complexity. Resulting parameters are presented in *F*.

From Table 3, it can be seen that the best fit model operates in the DDD regime (F=0.01), with demand-dependent detachment playing the major role in protein distribution. mProp is 0.49, which means that PSD95 is equally distributed between mobile and detached pools at the start of the study. The predicted degradation rate of PSD95 is higher than that for CA1. Therefore, the model predicted half-life is shorter (1.7 days against 4.3 days in CA1). This matches well with the observation of Bulovaite and colleagues (2022) for faster degradation in granule cells.

**Table 3.** List of parameters obtained in the optimisation for 3-month, 3-week and 18-month old mice in DG neurons. Bold indicates most striking changes between models.

| Parameter name | Definition | Value for 3M | Value for 3W | Value for 18M |
| --- | --- | --- | --- | --- |
| F | Ratio between DDT (1) and DDD (0) modes | 0.01 | 0.01 | 0.01 |
| Ctau | Coefficient for demand-dependent detachment | 2.3e-06 | 2.3e-06 | 2.3e-06 |
| mProp | Ratio between mobile and detached pools protein pools | 0.49 | 0.49 | 0.49 |
| demand_rd1 | Local demand value for rd1 | 1.03e-07 | 1.03e-07 | 1.03e-07 |
| demand_rd2 | Local demand value for rd2 | 0.029 | 0.029 | 0.029 |
| demand_rd3 | Local demand value for rd3 | 0.012 | 0.012 | 0.012 |
| demand_rd4 | Local demand value for rd4 | 0.041 | 0.041 | 0.041 |
| demand_rd5 | Local demand value for rd5 | 0.067 | 0.067 | 0.067 |
| demand_rd6 | Local demand value for rd6 | 0.089 | 0.089 | 0.089 |
| demand_rd7 | Local demand value for rd7 | 0.151 | 0.151 | 0.151 |
| demand_rd8 | Local demand value for rd8 | 0.610 | 0.610 | 0.610 |
| demand_rd9 | Local demand value for rd9 | 0.543 | 0.543 | 0.543 |
| demand_rd10 | Local demand value for rd10 | 0.281 | 0.281 | 0.281 |
| dv | Degradation rate (1/s) | 4.6e-06 | **5.52e-04** | **4.53e-06** |
| $T_{1/2}$ | PSD95 protein half-life, days | 1.74 | **0.014** | **1.77** |

We then tested if we were to predict the distribution of puncta density in 3-week and 18-month old mice simply by adjusting the degradation rates. As in the CA1 case, we fixed all model parameters except the degradation rates and refitted the model to the respective experimental data. Figure 5, C and D show the fit of the 3-month model to 3-week data without (*cost* = 0.014) and with (*cost* = 3.94e-16) adjusted degradation rates, respectively. Figure 5, E and F show the fit of the model to the 18-month data, again without (*cost* = 2.98e-05) and with (*cost* = 1.9e-09) adjusted degradation rates, respectively. As with the CA1 PyN model, we found that rate of degradation is much faster (*i.e*. shorter protein half-life) in 3-week old mice (varying in a range of hours instead of days), and only slightly longer in old mice compared to adult 3-month old mice (Table 3).

These results demonstrate consistency of the model effectiveness to explain the protein distribution in different types of neuron cell.

## Discussion

How developmental biology and genetics combine to build an organ as complex as the mammalian brain has been a major focus of neuroscience for many decades. The sheer scale and complexity of neuronal networks in brains is multiplied by the cellular diversity of neurons and molecular diversity of synapses that underpin each of these connections. The recent discovery of the hidden complexity of the synaptome architecture of the brain, where synapses of diverse shapes and molecular compositions are not distributed uniformly along the neuron and different synaptic subtypes can be observed at different localisations of the brain^7^, added a new dimension to the conceptual challenge of building and maintaining a complex brain. Although neuronal architecture is guided by a series of molecular genetic processes, it is highly unlikely that the spatial organisation of individual synapses is encoded by genetic programs that determine the molecular composition and dynamics of each synapse. Rather, a few developmental programs could build the synaptome architecture within the dendritic tree of a neuron and this architecture is then modified by neuronal activity and molecular interactions from the presynaptic terminal of input neurons.

Several computational models have previously attempted to describe the dynamics of molecular distribution in neurons (reviewed in ^11^). Models that assume diffusion as the prominent mode of molecular motion are not sufficient to describe experimentally observed protein distributions in dendritic spines, especially when considering the branched neuron structure^19–21^. This shortcoming can be addressed by including active transport of molecules along microtubules, which serve as railroads to provide uni- or bidirectional transport of both molecules and organelles along the long distances in axon and dendrites at a relatively fast velocity^22^. A conceptual model for active transport based on a “tug-of-war” between opposing forces of bi-directional motion was suggested by ^23^ and ^17^. Of particular interest for us was the sushi-belt model of demand-driven movement of molecules ^12,13,20,24^, which resolved the protein distribution with the minimal number of parameters for the whole realistic heavily-branched neuron morphology. The original sushi-belt model suggested by Doyle and Kiebler in 2011^12^ was designed to explain the conditional distribution of RNA and its supportive protein complex (RNPs)^25^. The model can be applied to other cargo including, synaptic vesicles, mitochondria, pigment granules, liquid droplets, viruses and intermediate filaments^22^, and proteins, notably PDZ domain-containing proteins in general^26^ and the PSD95 protein itself in particular^27^.

Here, we tested a tractable model that aims at describing a molecular basis for creating and maintaining synaptic diversity based on observed data. We demonstrated that our implementation of the sushi-belt model is sufficient for explaining the spatiotemporal distribution of PSD95: It can accurately describe the changes observed in pulse-labelled PSD95 distribution over 7 days, and, importantly, it can accurately predict the distribution of the protein in young and old animals by simply adjusting the protein degradation rate. We also showed that the model can be successfully applied to different neuron types, showing examples for CA1 pyramidal neurons and DG granule cells. Although we have specified a model for PSD95 data, the code built around this is generally applicable to any protein/RNA data and can be used with minor modifications.

As was suggested in ^13^, the sushi-belt model can operate in different modes, balancing the contribution of traffic and detachment to achieve more rapid or accurate distribution. We observed, for example, that our model prefers a demand-dependent trafficking (DDT) regime in CA1 neurons and demand-dependent detachment (DDD) in DG granule cell neurons. This suggests that resulting balance of the demand-dependent processes is flexible and is regulated by geometry or the molecular genetics of the cell.

Local demand in individual postsynaptic terminals is a dominant factor in our simulations but how this is regulated across the dendritic tree to produce the complex gradients is unclear. One possibility is that the postsynaptic synapse-specific differences in local demand could be induced by inputs from presynaptic terminals. These input projections would likely have different presynaptic protein compositions^28^, which in turn influence the clustering and localisation of postsynaptic proteins via trans-synaptic adhesion proteins. For example, the short protein lifetime of PSD95 observed in the slm may arise because the postsynaptic adhesion proteins may not have PDZ binding domains capable of stabilising PSD95. The involvement of instructive presynaptic adhesion proteins suggests that postsynaptic cell-autonomous mechanisms involving the sushi-belt distributes proteins and local demand is regulated by the presynaptic partner, a non-cell-autonomous process. Together these two processes could contribute to the complex synaptome architecture of individual neurons. Varying the postsynaptic processes of demand and degradation could then result in the observed age-related changes. These issues could be addressed in future studies, along with experiments that define the protein lifetime in specific classes of individual neurons, including interneurons.

Another factor regulating local demand could be the geometry of the dendritic tree. Williams at al., 2016 ^13^ described so-called “bottlenecks”, sites with very low demand, where the trafficking rates tend towards zero. Indeed, in our data we observed two sites with very low demand, namely CA1so_5 and CA1slm_1. Demand could be modulated by the activity state of the local synaptic environment. For example, locations with particularly low synapse density or very low branch number passing through these areas could dictate the urgency of molecular requirements. In this case, the demand structure should be less stable and will change according to the active states of specific synapses. Exploration of this hypothesis would require activity induced datasets. The likely scenario is that both components (synaptic structure and activity) may influence the final protein distribution, with a balance specific for the respective developmental stage and neuronal cell type.

Our results offer an enticingly simple framework that accounts for synapse specific delivery of molecules without resorting to more complex models where specific proteins are selectively trafficked to specific synapses. Individual synapses can effectively place a local request for more of a specific molecule from a cell-wide delivery system: The global transport mechanisms would provide a pool of molecules to meet local demand. If the global pool is depleted, then the cell just needs to act through generic homeostatic mechanisms to top up the global pool to ‘normal’ levels This does not preclude synaptic events leading to messages being sent back to the nucleus (or locally) to refine gene expression or a role for dendritic protein synthesis.

The framework presented here may help explain how many different diseases result in changes in the synaptome architecture and ultimately in abnormal behaviour. Defects in active transport have been linked to almost every neurodegenerative disorder (e.g. ^29^). Moreover, mutations in genes regulating proteostasis often correlate with autism, intellectual disability and synaptic dysfunction^30^. Mutations in synaptic proteins themselves can influence the turnover of other synaptic proteins^15^. Taken together with our results this implies that any genetic or environmental insult to the active transport system and/or the mechanisms of demand and degradation would rapidly challenge the neuron’s ability to maintain its normal synaptome architecture. Understanding these issues could be facilitated by building on our current simulations models to ones containing diverse interconnected neurons in a realistic connectome architecture, such as the cortical column. Further, including simulation of synaptic electrophysiological properties, could lead to integrated models of synaptome^31^ and connectome architectures at the structural and functional levels.

## Methods

### Mathematical background of the model

Full details for mathematical background of the model can be found in Williams et al, 2016^13^. The key points are reproduced below:

As described in ^13^, the steady-state ratio of cargo (PSD95 protein in our model) in neighbouring compartments equals the ratio of the trafficking rate constants (Figure 1C).

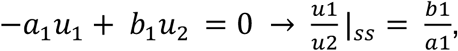

where *a* is forward rate constant (from the soma), *b* is backword rate constant (to the soma), u is concentration of PSD95 protein, attached to microtubules. Mass -action trafficking model can be re-expressed as a matrix differential equation, 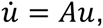 where *u* = {*u*_1_, *u*_2_, . . *u_N_*]^T^ is the state vector and A is the state-transition matrix. For a general branched morphology A will be nearly tridiagonal, with off-diagonal elements corresponding to branch points.

For three compartments the cable matrix will looks as follows:

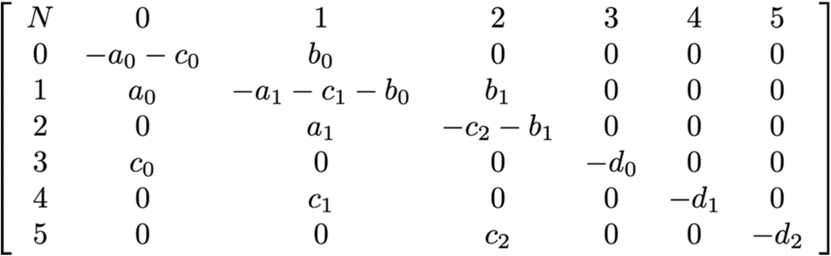

If detachment is included then for compartment *i* in a cable, the differential equation would take the form:

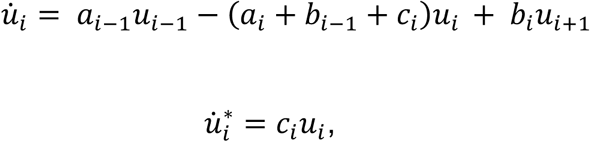

where *a* is forward rate constant, *b* is backword rate constant, c is rate of detachment, *u* is concentration of PSD95 attached to microtubules, and *u*^∗^ is concentration of the protein detached from microtubules (synaptic PSD in our case).

When *a_i_*, *b_i_* ≫ *c_i_*, then the distribution of cargo on the microtubules approaches the quasi-steady state as follows:

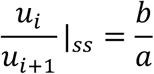

Following the ^13^ notation we introduced the local demand signal as a *ũ_i_*. From the previous equation we can derive conditions for protein concentration to match demand

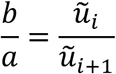

In this case protein will detach from microtubules with constant non-specific rate *c_i_* = *Const*, and the trafficking demand *ũ_i_*, which controls the distribution of the protein attached to the microtubules will be equivalent to the synaptic demand 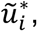 which controls the concentration of the free protein. Williams et al. called this regime Demand Dependent Traffic (DDT).

The alternative case called Demand Dependent Detachment (DDD) assumes that the trafficking rates are equals *a_i_* = *b_i_* and proteins are distributed evenly, and the synaptic demand 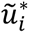 is fulfilled by demand-dependent detachment constant:

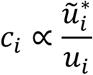

A whole spectrum of mixed modes that lies between DDT and DDD scenario is possible. For this, a scalar F, between 0 and 1, is introduced to describe mixed models. Assuming that the system is driven by the local synaptic demand *ũ*^∗^, which is normalised to sum to one, we need to choose *a_i_* and *b_i_* to satisfy the trafficking demand *ũ_i_* as follows:

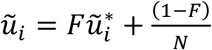

and *c_i_* to satisfy:

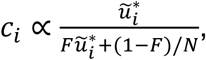

Where 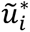 *i*s local synaptic demand in *i*th compartment, *ũ_i_* is local trafficking demand, *c_i_* is the local rate of detachment, *N* – number of compartments in the model and *F* is a ratio coefficient for DDT and DDD modes.

Setting F=1 results in DDT model (demand is satisfied by demand dependent trafficking only) and setting F=0 results in DDD model – demand is satisfied only by demand dependent detachment while trafficking rate constants remain uniform in all compartments.

### Model development

Initiating the data structures to support the neuron morphology model, preparation of the matrix of parameters and part of simulation code was adopted using Python code extracted directly from the Jupyter notebooks accompanying the ^13^ (https://github.com/ahwillia/Williams-etal-Synaptic-Transport). This code was then developed into Python package and extended to include the capacity for experimental data mapping and parameter extraction. All code is available on a GitHub repository (https://github.com/oksankas/Sushi_belt_PSD95).

### Model initialisation

Initial conditions of the system were assigned as follows: For each compartment, the normalized value 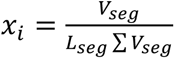 was calculated, where *V_seg_* is the experimental value for the segment or subregion at Day 0 and *L_seg_* is number of compartments in the segment or subregion. Then it was normalized to the total sum: 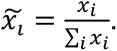 Cargo protein concentrations were assigned value 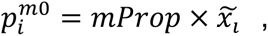 while synaptic protein concentrations were assigned value 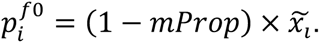

### Simulation and parameter optimisation

Parameter identification was performed using hybrid optimization techniques originally developed for the Systems Biology Software Infrastructure (SBSI)^32^. In short, 50 rounds of Particle Swarm Optimisation (PSO) were performed for a fast initial exploration of the parameter space, followed by 2000 rounds of Genetic Algorithm (GA) optimisation initiated using the best solutions found by the PSO step. Every 10 rounds of GA the best genome was optimized with 100 rounds of the Nelder-Mead algorithm before next generation construction to accelerate global optimum identification.

Population size for PSO was calculated as *N_PSO_* = *max*(15,5 × *N_par_*), where *N_par_* is the number of parameters to identify. Population for GA was selected as *N_GA_* = 2 × *N_par_*.

A cost function for the optimisation algorithm was calculated as the mean square difference between predicted concentration of the free protein pool at Day7 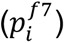 and normalized experimental value at that day 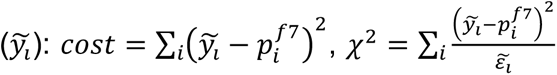 Normalized experimental values were calculated similarly to initial condition protein concentrations: 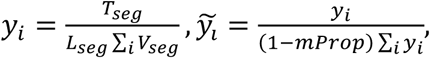 where *T_seg_* is the experimental value for the segment or subregion at the Day 7, normalized experimental error was calculated with relation to the Day 0 experimental value: 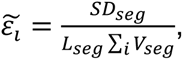 where *SD_seg_* is the experimental value standard deviation at the Day 7.

Note that normalisation of the Day 7 experimental value against the sum of Day 0 values requires us to take into account the decay of the total PSD95. Therefore, we used an additional normalisation factor (1 − *mProp*) because we are fitting only the synaptic component of the protein.

For the DG cell model each topological segment was divided into compartments of length 3.7 μm to be comparable with the CA1 model.

Neuron topology import and visualisation was performed using NEURON version 8.2.3 and an updated fork of PyNeuron-Toolbox (Alex H Williams. (2014) at *GitHub Repository*. https://github.com/lptolik/PyNeuron-Toolbox). Note that we selected Neuron to support future integration with electrophysiological models.

All calculations were performed on Edinburgh Compute and Data Facility (ECDF) the HPC cluster (https://digitalresearchservices.ed.ac.uk/resources/eddie) using python 3.10.12. All jobs used 8 compute cores with a total of 64 GB memory and a runtime limit of 48 hours.

## Acknowledgements

The authors would like to thank D. Maizels for artwork, C. Davey for editing and M. Kiebler for comments on the manuscript. For the purpose of open access, the authors has applied a CC-BY public copyright licence to any Author Accepted Manuscript version arising from this submission.

## Data availability

All code is available on a GitHub repository (https://github.com/oksankas/Sushi_belt_PSD95).

