## Supplementary Figures for "Protein trafficking and synaptic demand configure complex and dynamic synaptome architectures of individual neurons"

Supplementary Figure 1:

Model of a pyramidal CA1 neuron colour coded by values for normalised PSD95 puncta density taken on days 0 through day 7 of the Halo-ligand injection (from Bulovaite et al., 2022). See also main manuscript, figure 1A.

Supplementary Figure 2:

A. Distribution of the difference between a and b (a-b) rendered along the dendritic tree, shown along the normalised puncta density values at Day0 and Day7. It can be seen that a>b in most compartments and takes the max value in the most distal regions of the neuron (CA1lsms).

B. Same distribution as in A with the Demand values shown.

Supplementary Figure 3:

Distribution of average value for difference between *a* and *b* trafficking rate constants *(a-b)* for each of 20 segments of the dendritic tree as described in manuscript.

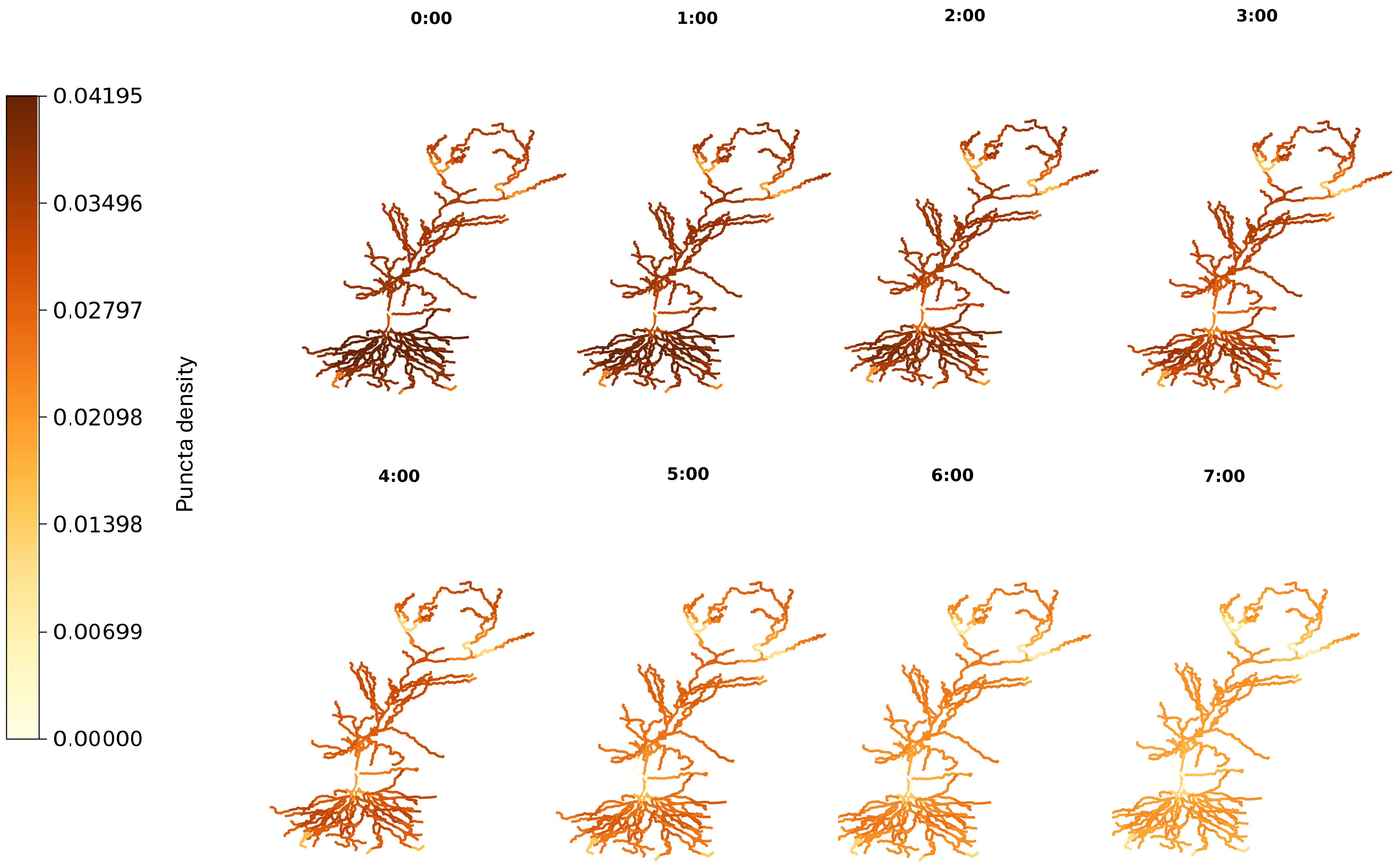
0.04195
0.03496
0.02797
0.02098
0.01398
0.00699
0.00000
Puncta density
0:00
1:00
2:00
3:00
4:00
5:00
6:00
7:00

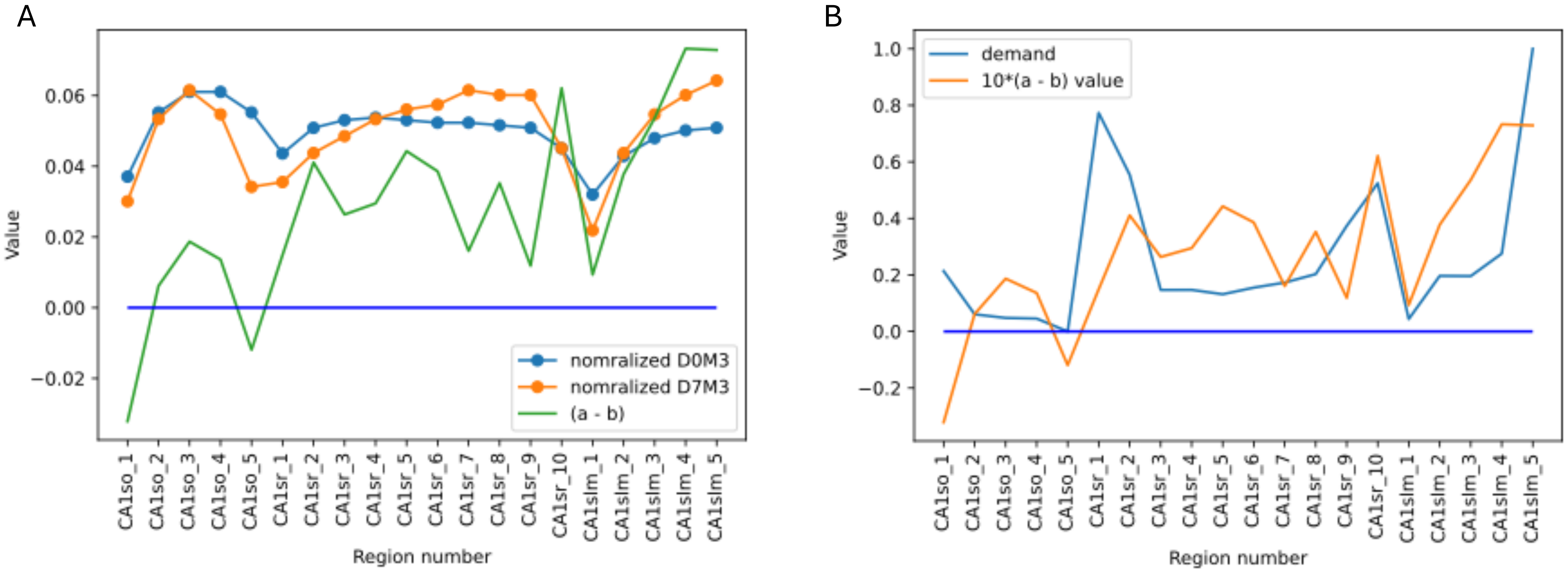
A
Value
nomralized D0M3
nomralized D7M3
(a - b)
Region number
B
demand
10*(a - b) value
Value
Region number
CA1so_1
CA1so_2
CA1so_3
CA1so_4
CA1so_5
CA1sr_1
CA1sr_2
CA1sr_3
CA1sr_4
CA1sr_5
CA1sr_6
CA1sr_7
CA1sr_8
CA1sr_9
CA1sr_10
CA1slm_1
CA1slm_2
CA1slm_3
CA1slm_4
CA1slm_5

### Average (a - b) distribution

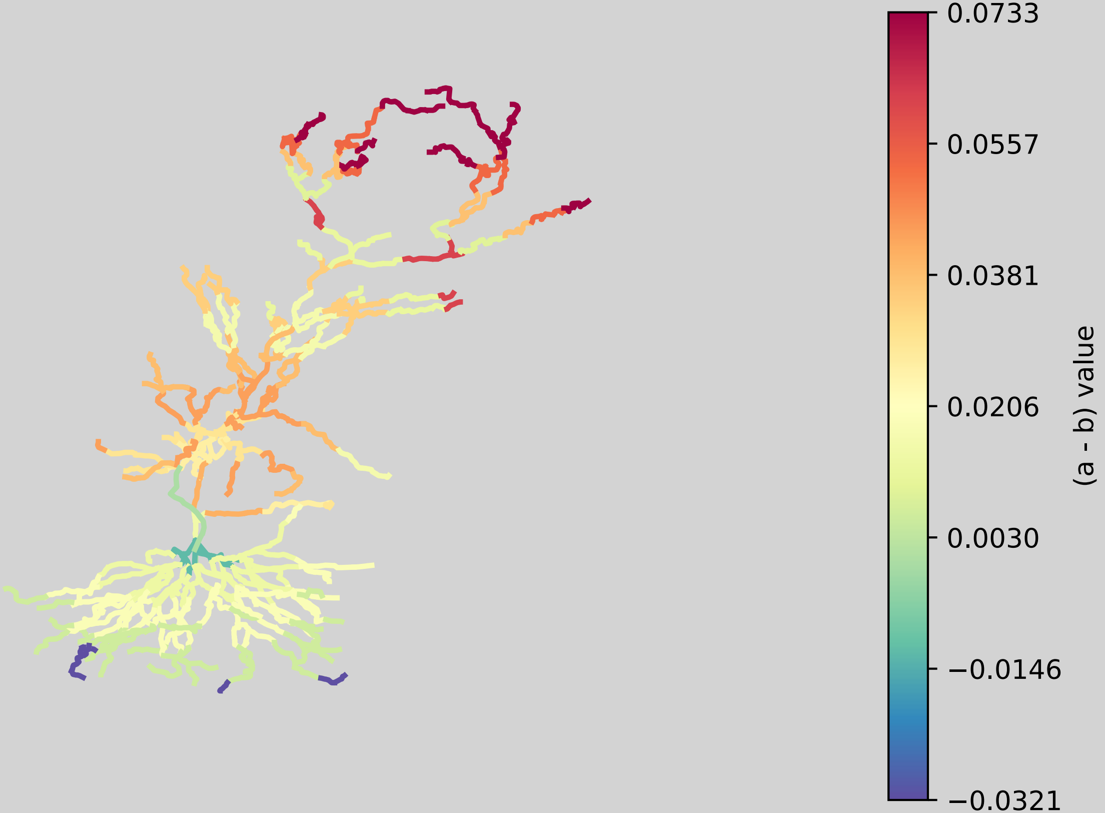
